# GWAS in Africans identifies novel lipids loci and demonstrates heterogenous association within Africa

**DOI:** 10.1101/2020.10.28.359497

**Authors:** Amy R. Bentley, Guanjie Chen, Ayo P. Doumatey, Daniel Shriner, Karlijn Meeks, Mateus H. Gouveia, Kenneth Ekoru, Jie Zhou, the Africa America Diabetes Mellitus Investigators, Adebowale Adeyemo, Charles N. Rotimi

## Abstract

**Background:** Serum lipids are biomarkers of cardiometabolic disease risk, and understanding the genomic factors contributing to their distribution has been of considerable interest. Large genome-wide association studies (GWAS) have identified over 150 lipids loci; however, GWAS of Africans (AF) are rare. Given the genomic diversity among those of African ancestry, it is expected that a GWAS in Africans could identify novel lipids loci. While GWAS have been conducted in African Americans (AA), such studies are not proxies for studies in continental Africans due to the drastically different environmental context. Therefore, we conducted a GWAS of 4,317 Africans enrolled in the Africa America Diabetes Mellitus study.

**Methods and Results:** We used linear mixed models of the inverse normal transformations of covariate-djusted residuals of high-density lipoprotein cholesterol (HDLC), low-density lipoprotein cholesterol (LDLC), total cholesterol (CHOL), triglycerides (TG), and TG/HDLC, with adjustment for three principal components and the random effect of relatedness. Replication of loci associated at p<5×10^−8^ was attempted in 9,542 AA. Meta-analysis of AF and AA was also conducted. We also conducted analyses that excluded the relatively small number of East Africans. We evaluated known lipids loci in Africans using both exact replication and “local” replication, which accounts for interethnic differences in linkage disequilibrium.

In our main analysis, we identified 23 novel associations in Africans. Of the 14 of these that were able to be tested in AA, two associations replicated (*GPNMB*-TG and *ENPP1*-TG). Two additional novel loci were discovered upon meta-analysis with AA (rs138282551-TG and *TLL2*-CHOL). Analyses considering only those with predominantly West African ancestry (Nigeria, Ghana, and AA) yielded new insights: *ORC5*-LDLC and chr20:60973327-CHOL.

**Conclusions:** While functional work will be useful to confirm and understand the biological mechanisms underlying these associations, this study demonstrates the utility of conducting large-scale genomic analyses in Africans for discovering novel loci. The functional significance of some of these loci in relation to lipids remains to be elucidated, yet some have known connections to lipids pathways. For instance, rs147706369 (intronic, *TLL2*) alters a regulatory motif for sterol regulatory element-binding proteins (SREBPs), which are a family of transcription factors that control the expression of a range of enzymes involved in cholesterol, fatty acid, and triglyceride synthesis.

## Introduction

Serum lipids are an important biomarker of cardiometabolic disease risk. As such, understanding the genomic factors contributing to their distribution has been of considerable interest. Large meta-analyses have identified over 150 loci associated with the distribution of serum lipids ^1–8^; GWAS in Africa are rare^9^. There are several reasons that this lack is concerning. First, as those of African ancestry have the greatest degree of genetic diversity in the world^10^, unless these populations are sufficiently represented in genomic research, a great deal of that diversity is not being evaluated for an association with serum lipids, potentially limiting the molecular insights that can be gained in our understanding of these traits. Second, although genome-wide studies of African Americans (AA) have been conducted, the drastically different environments of African Americans and continental Africans may differentially impact genetic associations. For instance, gene-environment interactions with environmental factors that differ between AF and AA have been found to affect serum lipid distribution, which may affect ability to detect a genetic association across populations^11–18^. As representation of those of African ancestry in genomic research has been relatively limited, it is common to analyze all those of African ancestry together to increase sample size; however, collapsing groups with such genomic and environmental diversity may obscure findings that are unique to populations within this broad category.

Here, we conducted a genome-wide association study of serum lipids in 4,317 Africans from the Africa America Diabetes Mellitus (AADM) study, a case-control study of type 2 diabetes (T2D) with sites in Nigeria, Ghana, and Kenya. Replication of GWAS-significant loci was sought in 9,542 AA. We report 23 novel loci that were genome-wide significant in Africans. Of the 14 of these that were available in the dataset of AA, two replicated. Novel loci were also identified when conducting analyses in West Africans separately, highlighting the importance of considering heterogeneity between African populations when conducting genomic analyses.

## Methods

### Participants

Participants included in this study were drawn from the Africa America Diabetes Mellitus (AADM) study, which has been previously described^19–21^. Briefly, AADM is a genetic epidemiology study of T2D, enrolling participants from university medical centers in Nigeria (Enugu, Lagos, and Ibadan), Ghana (Accra and Kumasi), and Kenya (Eldoret). Participants underwent a clinical examination that included a medical history, clinical anthropometry, blood pressure measurements and blood sampling.

Serum measurements were made on fasting samples. Serum lipids (triglycerides [TG], total cholesterol [CHOL], low-density lipoprotein cholesterol [LDLC], and high-density lipoprotein cholesterol [HDLC]) were determined enzymatically either with the COBAS Integra 400 Plus or Modular-E analyzers (Roche Diagnostics, Indianapolis, IN). Methods were standardized to in-house and other appropriate reference methods (e.g., CDC reference methods for HDLC, isotope dilution mass spectrometry for CHOL and TG). Missing LDLC measurements were calculated using the Friedewald equation^22^. Ethical approval was obtained by the National Institutes of Health and from the ethical committees in each study site. All participants gave written informed consent before participation in the study.

### Genotyping and Imputation

Genotyping was conducted either the Affymetrix Axiom^®^ PANAFR SNP array or the Illumina Consortium Multi-Ethnic Global Array (MEGA). Quality control was conducted separately for each of the resulting datasets. After technical quality control, sample-level genotype call rate was at least 0.95 for all subjects. Each SNP dataset was filtered for missingness (< 0.05), Hardy-Weinberg equilibrium (HWE, p < 10^−6^) and minor allele frequency (MAF > 0.01). SNPs that passed quality control filters were used as the basis for imputation. Imputation of all samples was done using the African Genome Resources Haplotype Reference Panel using the Sanger Imputation Service (https://imputation.sanger.ac.uk/). Variants were included in the analysis if they had a MAF ≥ 0.01 and an info score ≥ 0.3. All models were run using the dosage coding of variants.

### Statistical Analysis

Analyses were conducted using linear mixed models of the inverse normal transformations of lipids traits using EPACTS (version 3.2.6; http://genome.sph.umich.edu/wiki/EPACTS). Models were adjusted for age, age^2^, and sex. Additionally, we included the first three PCs of the genotypes, based on prior work^21,23^ (see **figure S4** of ^24^). Models were run with and without adjustment for body mass index (BMI), as an effect on lipids can occur through and independent of body adiposity. As this study is nested within a case-control study of T2D, models were also run with and without adjustment for T2D to evaluate a potential effect of disease status. All models also included a genetic relationship matrix to account for the random effect of relatedness, as related individuals were included in AADM. The genome-wide statistical significance level α for the main analysis was set at 5×10^−8^.

### Replication in African Americans

We sought replication for our genome-wide statistically significant associations in a combined dataset of African Americans (AA; n=9,542). Included AA samples were drawn from five datasets: the Howard University Family Study (HUFS)^25^ along with data available through dbGaP for the Atherosclerosis Risk in Communities study (ARIC phs000280.v2.p1, phs000090.v2.p1), the Cleveland Family Study (CFS phs000284.v1.p1), the Jackson Heart Study (JHS phs000286.v4.p1, phs000499.v2.p1), and the Multi-Ethnic Study of Atherosclerosis study (MESA phs000209.v13.p1, phs000420.v6.p3). When participants had more than one study visit, the visit with the most complete lipid measurements was selected, and if there were multiple visits with complete measurements, the most recent was selected (CFS). All studies obtained ethical approval from the relevant institutions and written informed consent from each participant prior to participation. Genome-wide genotyping data for all studies were imputed using the methods described above. Meta-analysis of AA study results was conducted with the inverse-variance-weighted fixed-effects method as implemented in METAL (https://genome.sph.umich.edu/wiki/METAL)^26^, and replication was assessed based on results of meta-analyses. Heterogeneity between East and West Africans was evaluated by with the p-value for heterogeneity using the “analyze heterogeneity” option in METAL.

### Analyses in West Africans

As our study included participants from both West Africa (Nigeria [n=2,207] and Ghana [n=1,376]) and East Africa (Kenya [n=734]), we investigated whether the inclusion of East Africans (EA) may have obscured findings that would have been detectable in WA alone. EA were not considered separately because of their relatively small sample size. Models in West Africans (WA) were run as described above for main analyses. Given the genetic similarity between AA and West Africans (WA), results from the WA analyses were meta-analyzed with AA data to take advantage of increased sample sizes for discovery.

### Replication of Known Lipids Loci

To evaluate the degree to which known lipids loci are also observed among Africans, we sought replication for 1,853 previously identified SNP-trait associations from large genome-wide studies of serum lipids ^1–8^. First, we conducted “exact” replication: evaluating the index variant reported in previous publications, with replication defined by consistent direction and p<0.05. Additionally, to account for differences in linkage disequilibrium (LD) between the initial reports (reflecting predominantly European ancestry) and our African samples, we conducted “local” replication. Local replication is an LD-based replication technique, investigating all variants with r^2^>0.3 with the index variant among CEU. A locus was included in the local replication analysis if there was no exact replication of that locus in our data and if there were additional variants with r^2^>0.3 [CEU] with the index variant that passed filtering criteria and were available in our data. Associations were considered replicated locally with consistent direction (based on direction of association of minor allele for index and local variant) and p < 0.05 / LD-based effective number of tests in the region (calculated as in ^27^).

## Results

The study population was heavier and had a worse lipid profile than would be expected of a general population sampled in these regions, consistent with the facts that this study was nested within a case-control study of T2D and approximately half of participants were T2D cases (Table 1). Additionally, all recruitment was conducted in urban areas, and urban location is associated with worse cardiometabolic outcomes in West Africa ^28^.

**Table 1.** Descriptive Statistics for Included Participants.

|  | *West Africans* | *East Africans* | *All* |
| --- | --- | --- | --- |
| N | 3583 | 734 | 4317 |
| Age (years) | 50.6 (13.2) | 55.4 (10.2) | 51.4 (12.9) |
| BMI (kg/m<sup>2</sup>) | 26.6 (5.7) | 26.4 (5.4) | 26.6 (5.6) |
| Type 2 Diabetes* | 50.7% | 50.7% | 50.7% |
| HDLC (mg/dl) | 45.3 (16.7) | 47.4 (16.8) | 45.6 (16.7) |
| LDLC (mg/dl) | 131.1 (46.6) | 130.0 (46.3) | 130.0 (46.3) |
| TG (mg/dl) | 98.3 (48.3) | 167.5 (91.9) | 110.0 (63.5) |
| CHOL (mg/dl) | 196.8 (52.9) | 206.2 (58.0) | 198.4 (53.9) |
| TG/HDLC | 2.63 (2.13) | 3.96 (2.77) | 2.85 (2.3) |

*Mean (SD) reported, except where indicated. \*Proportion shown.*
*BMI: body Mass Index; TG: triglycerides; HDLC: high density lipoprotein cholesterol; LDLC: Low density lipoprotein cholesterol; CHOL: total cholesterol*

Our main GWAS analysis in AADM identified 23 novel GWAS-significant lipids loci (29 variants) in AADM (**Table 2; **Supplementary Figures 1** [Manhattan plots] and 2 [QQ plots]**), with novelty defined by distance from known lipids loci (> 1 MB) and statistical significance set at 5 × 10^−8^. Notable among these findings were associations of variants with higher frequency among Africans compared to other worldwide populations, perhaps allowing for their detection in this analysis. For instance, rs138202830 (*GBE1*), rs188701119 (*TINAG*), rs140987192 (*PBX3*), and rs75360819 (*CDH2*) are monomorphic in non-AFR 1000 Genomes populations, and the minor alleles for rs111590558 (*HTR2A*), rs7281821 (*APP*), and rs144086909 (*LARGE*) are at their highest frequencies in the AFR super-population (Table 2).

**Table 2.**
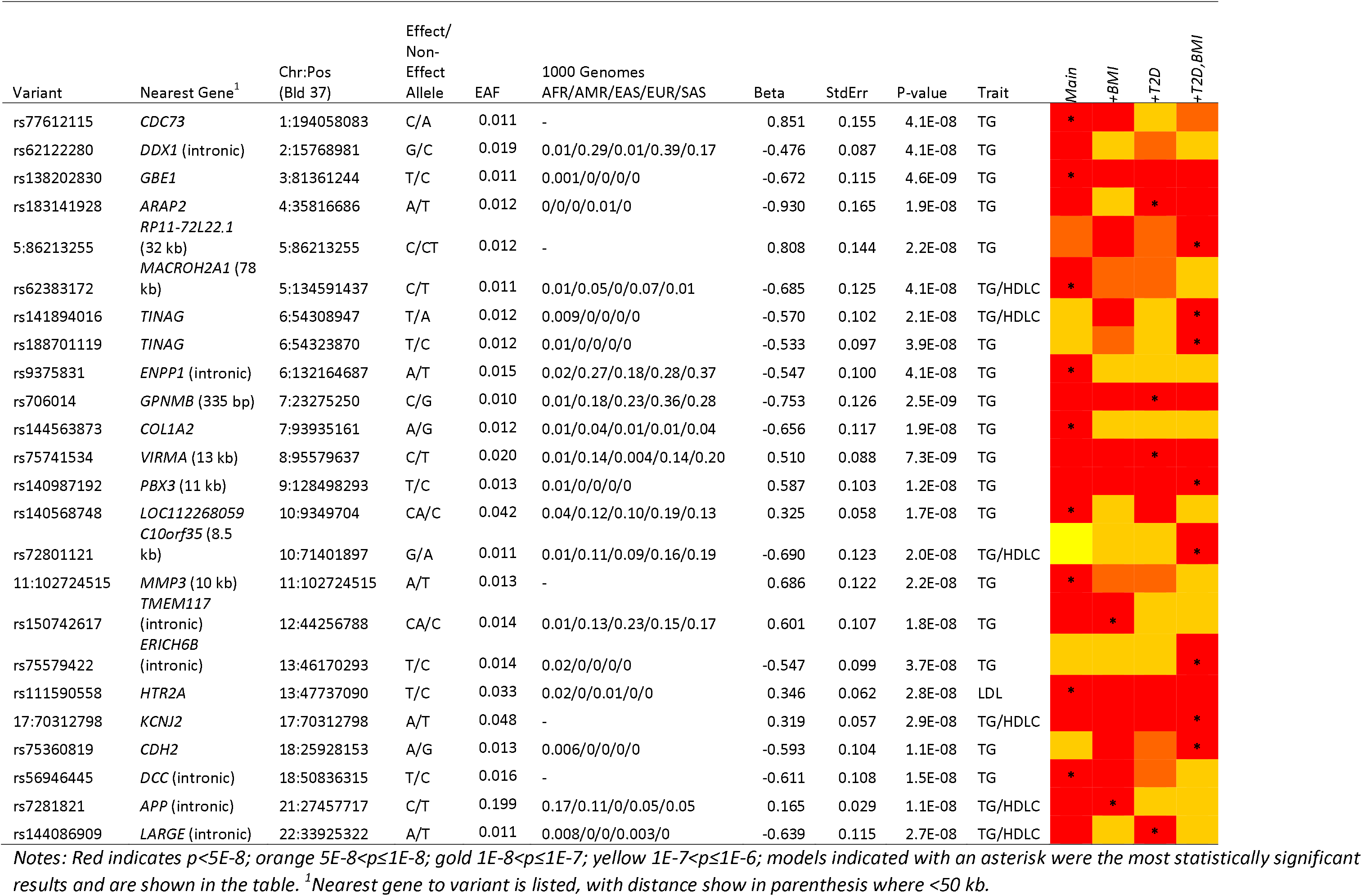
Genome-Wide Lipids Associations p<5E-08 among Africans (n=4,317)

**Figure 1.**
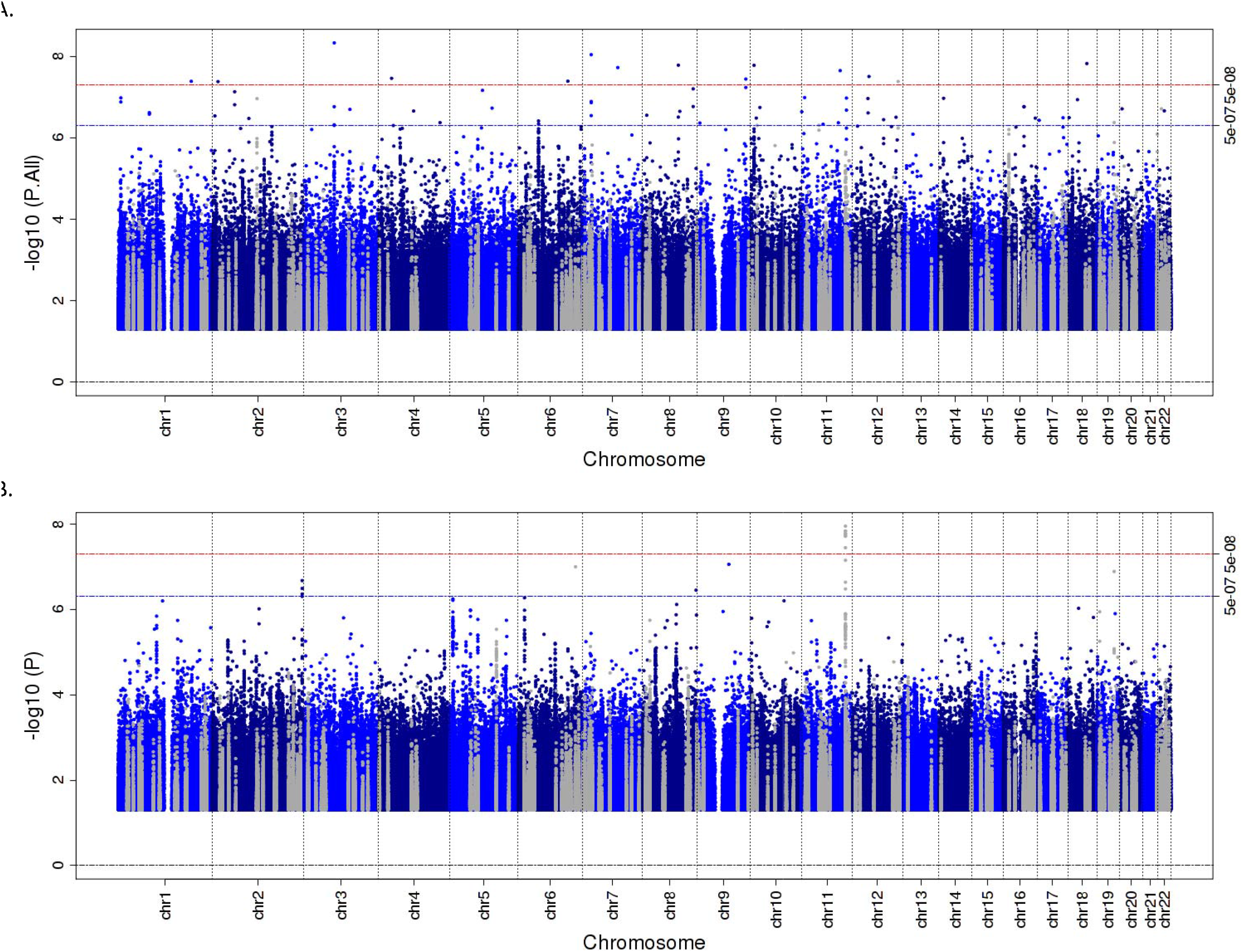
Manhattan Plots for TG in West and East Africans combined (A.) and in West Africans only (B.)

Given the potential impact of both BMI and T2D on associations with serum lipids, particularly in this dataset nested within a case-control study of T2D, we tested models with and without adjustment for these two covariates. In terms of statistical significance, adjustment for BMI and T2D had relatively minor effects. For 22 of the 23 novel loci, associations that were statistically significant in one model had p ≤ 10^−7^ for all other models (for the remaining locus, all models were p ≤ 10^−6^; Table 2).

To determine the degree to which these associations were also found among AA, we evaluated these loci in 9,542 AA (**Supplemental Table 1**). Of 29 variants in 23 loci investigated, 16 variants in 14 loci passed frequency and quality control criteria and were able to be evaluated (**Supplemental Table 2**). Two of these loci were replicated (p<0.05 with matching direction of association). The minor (G) allele of rs706014 (5’ of *GPNMB*, Glycoprotein Nonmetastatic Melanoma Protein B) was associated with lower TG in both Africans (β −0.75 TG, p=2.5 × 10^−9^) and in AA (β −0.084 TG, p=0.0093). The minor (T) allele of rs9375831, intronic to ENPP1 (ectonucleotide pyrophosphatase/phosphodiesterase 1), was also associated with lower TG in Africans (β −0.55 TG, p=4.1 × 10^−8^) and in AA (β −0.076 TG, p=0.037). As variants in this gene have been previously associated with insulin resistance and related traits^29–34^, we also investigated this relationship in our data. An association of the minor allele of rs9375831 with HOMA-IR in AF participants without T2D did not reach statistical significance (p=0.2), but was directionally consistent: the minor allele was associated with decreased TG and decreased HOMA-IR. We also evaluated the ratio of TG to HDL, a biomarker of insulin resistance: the minor allele was associated with decreased TG/HDL (p=0.0002), though this association may be largely driven by the association of this variant with TG.

We conducted meta-analyses of Africans and AA, which yielded two additional GWAS-significant loci (**Table 3**). The minor (T) allele of intergenic variant rs138282551 was associated with higher TG in both Africans (β 0.29 TG, p=4.5 × 10^−7^) and in AA (β 0.14 TG, p=0.0035). This variant is at much higher frequency among those of African ancestry compared to other populations (among 1000 Genomes super-populations: AFR 0.03, AMR 0, EAS 0, EUR 0.002, and SAS 0). The minor (T) allele for an African ancestry-specific intronic variant in *TLL2*, rs147706369, was associated with higher total cholesterol, with similar magnitude (β 0.34 CHOL [AFR], β 0.28 CHOL [AA]) and strength (p=1.5 × 10^−4^ [AFR] and p=2.6 × 10^−5^ [AA]) of association for both groups.

**Table 3.** Novel Signals in Meta-Analysis of Africans and African Americans.

| rsID | Chr:Pos<sup>1</sup> | Eff/Non-Eff<br>Alleles | 1000 Genomes MAF<br>AFR/AMR/EAS/EUR/SAS | Trait<sup>2</sup> | Analysis | N | EAF | Beta | SE | P-Value |
| --- | --- | --- | --- | --- | --- | --- | --- | --- | --- | --- |
| rs138282551<br>*Intergenic* | 1:81649749 | T/C | 0.03/0/0/0.002/0 | TG | Africans | 4292 | 0.04 | 0.29 | 0.058 | 4.5E-07 |
|  |  |  |  |  | Afr. Americans | 8537 | 0.03 | 0.14 | 0.049 | 3.5E-03 |
|  |  |  |  |  | Meta-analysis | 12829 | 0.035 | 0.20 | 0.037 | **4.6E-08** |
| rs147706369<br>*TLL2*<br>*Intronic* | 10:98213172 | T/C | 0.016/0/0/0/0 | CHOL | Africans | 4300 | 0.016 | 0.34 | 0.090 | 1.5E-04 |
|  |  |  |  |  | Afr. Americans | 9217 | 0.015 | 0.28 | 0.067 | 2.6E-05 |
|  |  |  |  |  | Meta-analysis | 13517 | 0.016 | 0.30 | 0.054 | **1.7E-08** |

<sup>1</sup>*Bld. 37;* <sup>2</sup>*All traits inverse normal transformed.*

Our main analyses of Africans included participants from both West Africa (Nigeria [n=2,207] and Ghana [n=1,376]) and East Africa (Kenya [n=734]). Given the genetic and environmental differences among the sampled populations, it is possible that a signal present in participants from only one of these geographic regions could be masked when the two were included together. Additionally, mean values for TG and closely related values (TG/HDLC and CHOL) differed dramatically in East compared to West Africans (TG 167.5 vs. 98.3 mg/dl, respectively; **Supplemental Table 3**). Based on these observations, we conducted a separate analysis of WA; a separate analysis of EA is not presented due to the relatively small sample size. There were no novel statistically significant hits when WA were considered on their own, but known signals that were not apparent in the full GWAS were observed in the analysis of WA only, including the known chromosome 11 locus *BUD13* and, to a lesser extent, *APOE/APOC1* on chromosome 19 (**Figure 1**). As AA have predominantly West African ancestry (79.9%^35^), meta-analysis of WA and AA was also conducted, and two additional hits were identified (**Table 4**). The minor (C) allele of rs7797481 (*ORC5*) was associated with higher LDLC in both WA and AA. The association was quite different among the EA (β −0.38, p=0.053; p for heterogeneity between WA and EA=0.0028), and their inclusion in the meta-analysis reduced the statistical significance of the association (**Figure 2a**). In contrast, for the association between the minor (T) allele of 20:60973327 (chromosome:position [build 37]) and higher TG in the meta-analysis of WA and AA, the effect size was similar among EA but with a large standard error (β 0.13, SE=0.11), making the association above our threshold for statistical significance when they were included in the meta-analysis (**Figure 2b**; p for heterogeneity between WA and EA=0.50).

**Table 4.** Novel Signals in Meta-Analysis of West Africans and African Americans.

| rsID | Chr:Pos<sup>1</sup> | Eff/Non-Eff Alleles | 1000 Genomes MAF AFR/AMR/EAS/EUR/SAS | Trait<sup>2</sup> | Analysis | N | EAF | Beta | SE | P-Value |
| --- | --- | --- | --- | --- | --- | --- | --- | --- | --- | --- |
| rs7797481<br>*ORC5*<br>intronic | 7:103770056 | C/G | 0.02/0.13/0/0.20/0.07 | LDLC | West Africans | 3566 | 0.028 | 0.25 | 0.075 | 8.4E-04 |
|  |  |  |  |  | Afr. Americans | 8501 | 0.057 | 0.16 | 0.035 | 2.5E-06 |
|  |  |  |  |  | Meta-analysis | 12067 | 0.052 | 0.18 | 0.031 | **1.4E-08** |
| - | 20:60973327 | T/C | -- | CHOL | West Africans | 3565 | 0.110 | 0.21 | 0.049 | 2.2E-05 |
|  |  |  |  |  | Afr. Americans | 8901 | 0.105 | 0.13 | 0.035 | 2.2E-04 |
|  |  |  |  |  | Meta-analysis | 12466 | 0.107 | 0.16 | 0.029 | **4.4E-08** |

<sup>1</sup>Bld. 37; <sup>2</sup>All traits inverse normal transformed.

**Figure 2.**
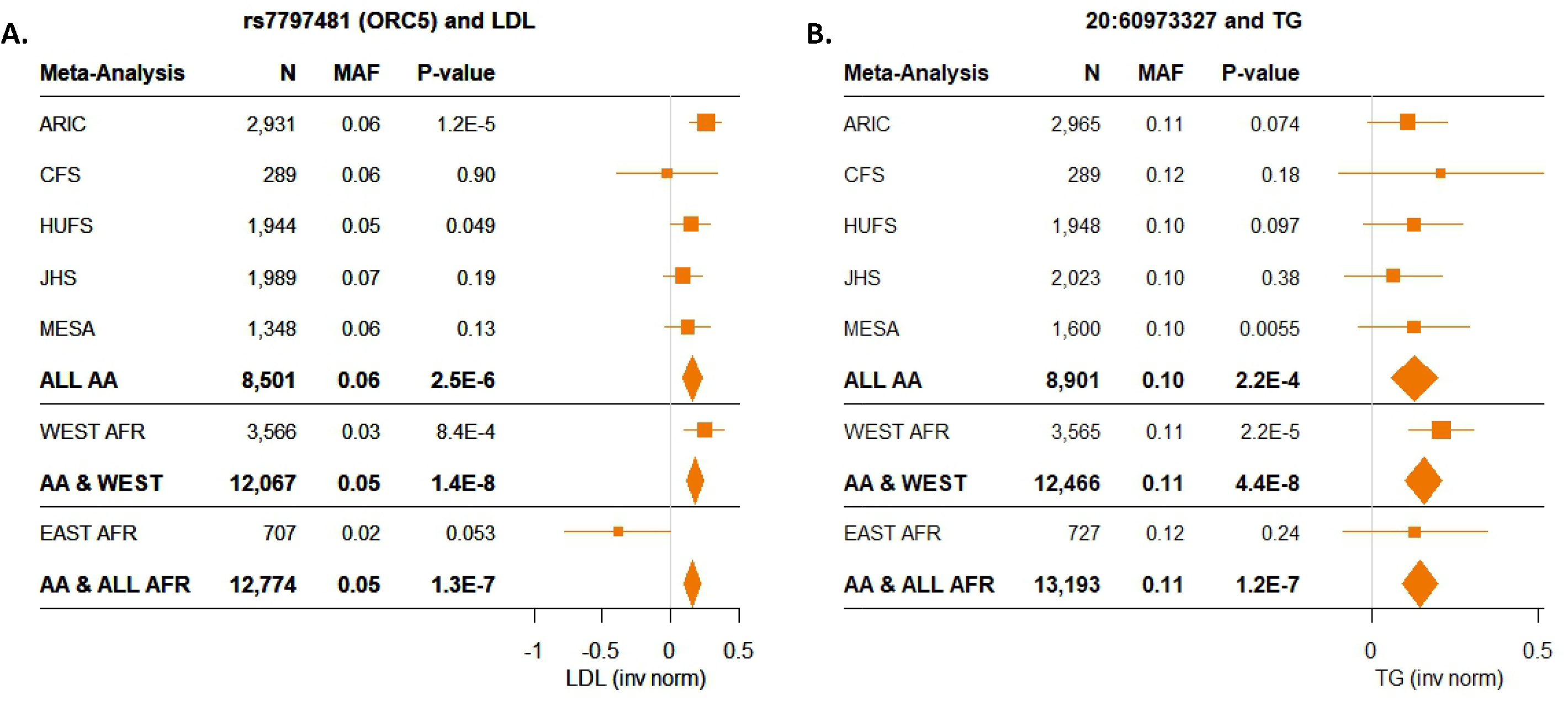
Novel Associations in Meta-Analysis of West Africans and African Americans

We sought replication for 1,853 previously identified SNP-trait associations from large genome-wide studies of serum lipids ^1–8^. Of 1,534 SNP-trait associations available to be tested in our data, 947 were in a consistent direction (62%), and 144 of those replicated (consistent direction and p<0.05). This represents an exact replication rate of 9.4% (**Supplemental Table 4**). Of the SNP-trait associations that did not replicate in the exact analysis, 702 had variants that were suitable for evaluation using “local” replication (an LD-based replication technique, see Methods). An additional 27 SNP-trait associations in 24 loci were replicated using these criteria (**Supplemental Table 5**). In total, we replicated 83 out of 303 loci (27%).

## Discussion

In this GWAS of serum lipids in 4,317 continental Africans, we identified 23 novel lipids loci. Of these, 14 loci were available to be tested in 9,542 African Americans for replication and two of these associations replicated (*GPNMB*-TG and *ENPP1*-TG). Upon meta-analysis of the Africans and African Americans, two additional associations were found (rs138282551-TG and *TLL2*-CHOL). Given genetic and phenotypic diversity between WA and EA, we also conducted analyses excluding EA. Two additional novel associations were identified with meta-analysis of WA and AA: *ORC5*-LDLC and chr20:60973327-CHOL.

Our focus on African populations may have allowed for novel TG findings based on both the understudied environmental and genomic context of these individuals. The vast majority of our findings were for triglycerides. In a recent study assessing transferability of known lipids loci from European ancestry populations to Ugandans, approximately 70% of HDLC and LDLC loci were transferable to Ugandans, whereas only 10.5% of TG loci were transferable. The authors suggested that this difference may have resulted from differences in the environmental context^36^. While environmental context is a likely contributor to variation in TG levels, differences in genetic context may also be important.

Descriptive data from a recent serum lipids meta-analysis of nearly 100 cohorts under the aegis of the CHARGE consortium offers useful comparisons in terms of ancestry-level differences in TG ^16^. Mean TG levels in AA are relatively low compared to those of other ancestries (~ 100 mg/dl for all included AA studies, except ARIC [115 mg/dl]) and similar to WA (mean 98.3 mg/dl), despite dramatically different environmental backgrounds. In contrast, the mean values for EA were markedly higher: mean 168 mg/dl. This distribution suggests that genomic factors underlie some of the variation across populations in TG levels.

As the numbers of studies and samples with African ancestry are generally low relative to European ancestry, studies including individuals of African ancestry are generally considered as a single group. However, this grouping may not be appropriate given the genetic diversity among those who could be categorized as having African ancestry and the diverse environmental backgrounds in which those of African ancestry reside. Given that our study population included individuals from both West and East Africa, who are distinct in terms of genetics and diet, we conducted analyses separately for WA. The exclusion of EA from the analyses, leaving the two groups with predominantly West African ancestry, allowed for the discovery of novel loci, presumably because of genetic similarities between the WA and AA, in contrast to the EA. The AA are still not an ideal replication sample for the WA, however, given both their admixture (with an average of 20% European ancestry) and the environmental background, which is quite distinct between those of African ancestry living in West Africa and in the United States. Thus, the differences in observed associations between these groups are likely to represent a combination of factors, including genetic differences, environmental context, and false positives, in unknown proportions.

One emerging potential explanation for differences by ancestry in associations with lipids is through an effect of diet on the gut microbiome^37^. Gut microbiome features have been shown to underlie some of the interindividual variation in lipids-related protein levels^38^. Transplant of microbiota from humans with dyslipidemia to germ-free mice led to a recapitulation of the lipid phenotype^39^. Diet differs for each of the groups included in this analysis^40–42^, and this could contribute to differences in the associations observed. Further analyses including individuals of varying degrees of genetic and environmental differences, particularly those including microbiome data, will help to advance our understanding of these relationships in the context of serum lipids.

Higher BMI and T2D are both associated with dyslipidemia, and Mendelian randomization studies suggest that BMI and T2D are causal for dyslipidemia^43,44^. As such, we sought to establish whether associations were independent of an effect on either of these traits by conducting all analyses with and without adjustment for BMI and T2D. Given that this study is nested within a case-control study for T2D, and, thus, participants had higher proportions of T2D and elevated BMI compared to what would be expected in a general population study, it was of particular importance to account for the potential impact of these factors on our analyses. In our experience, however, adjusting for either of these factors did not alter the findings considerably. For all of our main findings, statistical significance of the associations was quite similar across models, although some of the minor fluctuations meant that some associations in some models met our criteria for statistical significance, while others just missed. For all of our 23 lead associations, all other models yielded associations with p ≤ 10^−6^, and for 22 of them, all other models yielded association with p ≤ 10^−7^. Based on these findings, while the presence or absence of BMI or T2D in the model may have made some signals easier to detect, they did not wipe out any associations, as would have been expected if an association with lipids was occurring through an effect on BMI or T2D.

The proportion of known lipids loci that replicated among Africans was 27%. Attempted replication of the previously reported index SNPs yielded a replication rate of only 9.8%. This was increased to 27% by using a linkage disequilibrium-based “local” replication method, which accounted for the generally lower linkage disequilibrium among African ancestry individuals by evaluating all of the variants in LD with the index variant in the previously reported population (with correction for the number of variants tested). These proportions are similar to those reported for replication of known type 2 diabetes loci in Africans, in which 15% replicated exactly and an additional 15% replicated using a local replication method^21^, giving us confidence that these findings are in a reasonable range. While this relatively low proportion reflects in part the smaller sample size present for the Africans compared to the very large meta-analyses in which many of these loci were identified, there are other factors that may have contributed. For instance, there is evidence of differential associations across ancestry^45^. Additionally, the environmental background of the Africans compared to those more well-represented populations could play a role. A recent analysis of the transferability of lipids loci to Ugandans reported that among major lipids loci (p<10^−100^ in the largest European ancestry meta-analysis), 71% of those that were not transferable to Ugandans showed evidence of pleiotropy with BMI among European ancestry populations, while none of those that were transferable were also associated with BMI (these mostly represented well-established lipids pathways). The authors theorize that these discrepancies may stem from gene-environment interactions, with effects on lipids not shared across different environmental backgrounds. For these major known lipids loci, evidence for and against transferability to our West and East African study populations matched very closely the results in the study in Ugandans (90% concordance)^36^.

In our analysis, the A allele for rs9375831 (intron, *ENPP1*) was associated with lower TG in Africans, and this association was replicated in AA. This variant is in promoter and enhancer histone marks for liver tissue (Haploreg^46^: https://pubs.broadinstitute.org/mammals/haploreg/haploreg.php). This gene is known for a role in the development of insulin resistance, of which hypertriglyceridemia is a hallmark. *ENPP1*, also known as PC-1, directly interacts with the α subunit of the insulin receptor, inhibiting insulin signaling^47^. Transgenic mouse models of overexpression of *ENPP1* in muscle and liver^48^ and adipose tissue^49^ develop insulin resistance, including hypertriglyceridemia^49^. It has been hypothesized that higher *ENPP1* expression leads to reduced triglyceride storage capacity, increased fat deposition in the liver, and subsequent systemic abnormalities associated with insulin resistance^49^. Increased expression of *ENPP1* in adipose tissue of non-obese, non-diabetic individuals is associated with a reduction in insulin sensitivity^50^. Variants in this gene (not in LD with rs9375831) have been associated with insulin resistance and related traits in multiple analyses^29–34^, with evidence for elevated triglycerides reported for some^29,34^, but not all studies ^30–33^.

We found that the minor (C) allele for rs706014 (335 bp 5’of *GPPMB*) was associated with decreased serum triglycerides. According to Gene-Tissue Expression project (GTEx) data, this variant has 44 expression quantitative trait loci (eQTL) hits in a variety of tissues. In adipose tissue, this variant is associated with circulating levels of *GPNMB*, *NUPL2*, and *KLHL7-AS1*^46^. GPNMB is involved in pro-inflammatory responses and tissue repair. Animals fed high fat diets experienced large increases in Gpnmb expression in ascending aorta of rabbits (15-fold)^51^ and visceral adipose tissue of mice (30-fold change)^52^. A lysosomal storage disease involving intracellular accumulation of unesterified cholesterol, Niemann-Pick disease, is also associated with elevated levels of Gpnmb in tissue and plasma in both model organisms and in humans, with levels increasing with age and disease progression ^51^. The minor alleles of two variants in moderate LD with rs706014 (rs199351 and rs199347, R^2^ ~0.3 among EUR), which are also eQTLs for GPNMB (Haploreg), have been associated with decreased risk of Parkinson’s Disease (PD)^53–55^. Higher lipid levels, including specifically triglycerides^56^, have been associated with decreased PD risk ^57,58^. Emerging evidence of a complex interplay between lipids and α-synuclein, a protein that is strongly associated with PD, in disease risk has even led some researchers to suggest that PD should be considered a lipidopathy^59^.

We report that the minor allele (T) for rs138282551 (intergenic) is associated with higher TG among Africans and AA. This variant alters motifs for CHD2, E2F, and TCF4^60^. Studies of E2F1 knockout mice demonstrated that E2F1 is involved in global transcriptional regulation of de novo lipid synthesis. In mice, E2F1 expression and activity in the liver increased in response to feeding and insulin levels, and E2F1 has been characterized as a major regulator of lipid metabolism^61^. Deletion of *E2f1* in cellular and mouse models leads to decreased Pcsk9 expression and increased LDLR expression, and abnormal cholesterol accumulation in the liver^62^.

In our meta-analysis of AFR and AA, the minor allele of rs147706369 (intronic, *TLL2*) was associated with increased total cholesterol. This variant alters a regulatory motif for sterol regulatory element-binding proteins (SREBPs)^60^, which are indirectly required for cholesterol biosynthesis. SREBPs are a family of transcription factors that control the expression of a range of enzymes involved in cholesterol, fatty acid, and triglyceride synthesis, and have been called the “Master Regulators of Lipid Homeostasis” ^63^.

The minor allele of rs7797481 (*ORC5*, intron) was associated with higher LDLC in WA and AA. This variant is an eQTL for *ORC5L* in whole blood^64^. The origin recognition complex (ORC) proteins play a central role in the initiation of DNA replication. It is unclear how this could be related to the current findings. This variant also alters regulatory motifs for FOXA and GR^60^. Members of the FOXA family have been shown to alter lipid metabolism through regulation of multiple lipids pathways. Foxa1 inhibits the expression of many of the genes involved in triglyceride synthesis and accumulation and in VLDL synthesis, including *APOB* (which encodes the major structural component of LDLC), in human hepatocytes^65^. Foxa1 also inhibits triglyceride synthesis and accumulation in human hepatocytes, and FOXA1 is downregulated in non-alcoholic fatty liver disease^66^.

Our conclusions should be evaluated in the context of our study’s strengths and weaknesses. One strength of this study is the large sample size for a study of continental Africans, who remain understudied in genomic analyses for serum lipids. Notably, all these individuals were recruited as part of one study, with identical questionnaires and assay methodology, to limit the potential for observed differences in East vs. West Africans being created by study methods. Differences among African ancestry individuals are explored by considering West and East Africans in stratified analyses rather than treating “African ancestry” as a monolithic category. The dataset of AA, all imputed together using the same reference panel, brought an additional large number of African ancestry samples to the analysis. We used both an exact and a local replication method for assessing transferability of previously identified loci to Africans thereby ensuring that we had accounted for the LD-based differences in the discovery and replication datasets. It would have been ideal to have replication cohorts that were a better match for the environmental and genomic background of the West Africans in our study for replication. With the replication cohorts that we used, we were unable to determine whether association differences represented false positives, differences in environment, or differences in genomic context. We look forward to future studies in which these types of analyses are possible.

In summary, we conducted a genome-wide association study of serum lipids in Africans in Nigeria, Ghana, and Kenya, with meta-analysis with and replication in AA. Among the overall study findings, several of the novel loci are important in TG or cholesterol synthesis, storage or turnover. While functional work will be useful to confirm and to understand the biological mechanisms underlying these associations, this study demonstrates the power of conducting large-scale genomic analyses in Africans.

## Acknowledgements

The authors would like to gratefully acknowledge the AADM collaborators, including Olufemi Fasanmade, Thomas Johnson, Johnnie Oli, Godfrey Okafor, Benjamin A. Eghan Jr., Kofi Agyenim-Boateng, Jokotade Adeleye, Williams Balogun, Clement Adebamowo, Albert Amoah, Joseph Acheampong, and Duncan Ngare (deceased). The views expressed in this manuscript are those of the authors and do not necessarily represent the views of the NIH.

