## Supplemental Figures for "GWAS in Africans identifies novel lipids loci and demonstrates heterogenous association within Africa"

Supplementary Figure 1: Manhattan Plots for Serum Lipids in AADM in a model adjusted for age, age^2^, sex, and 3 PCs.

A. Cholesterol

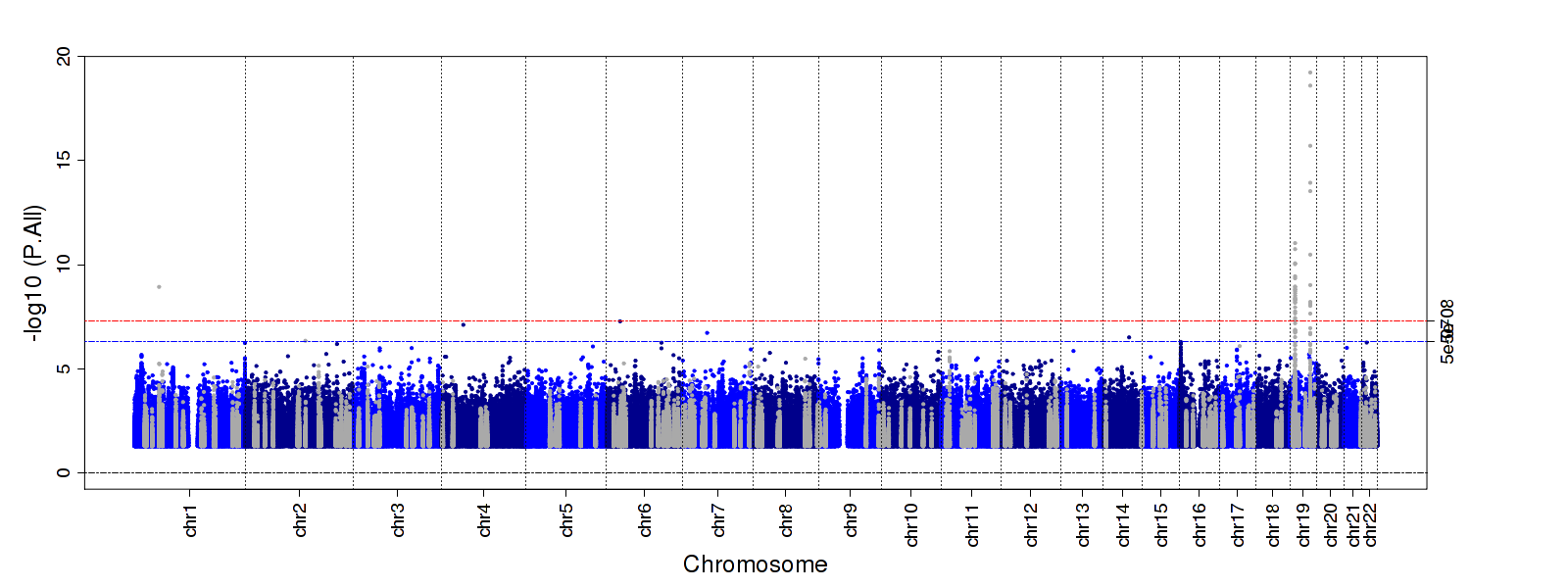

B. HDL

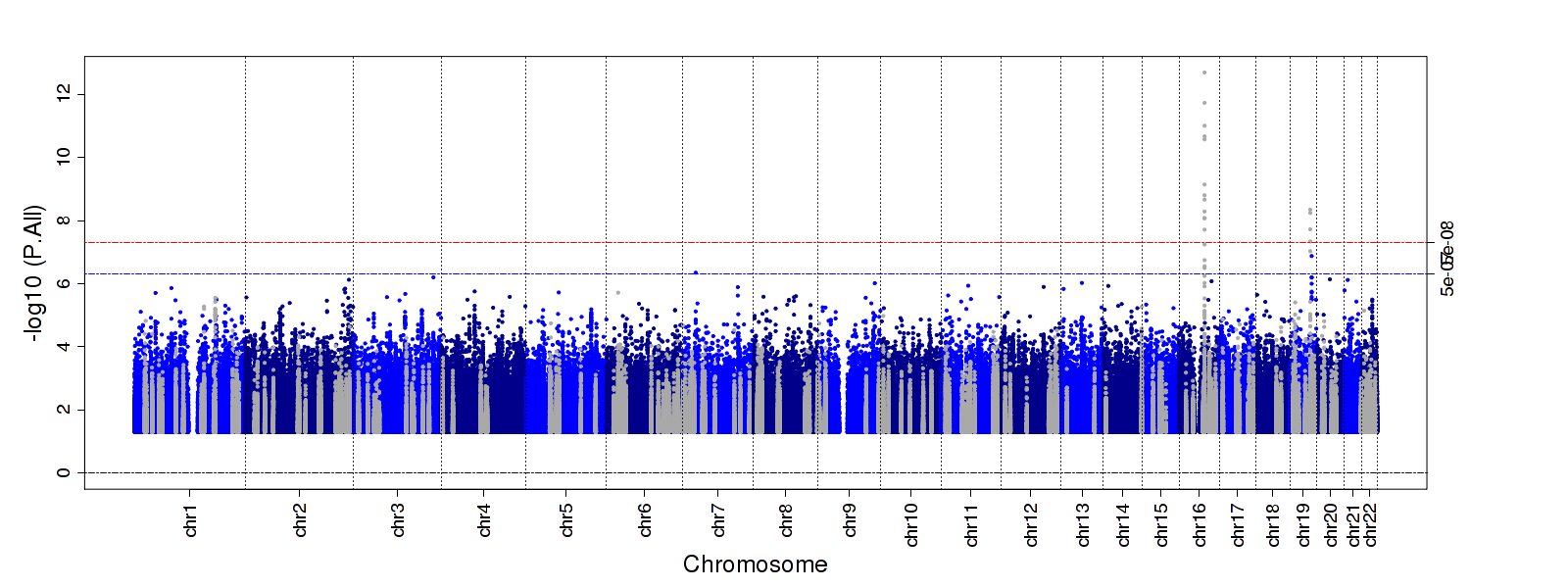

C. LDL

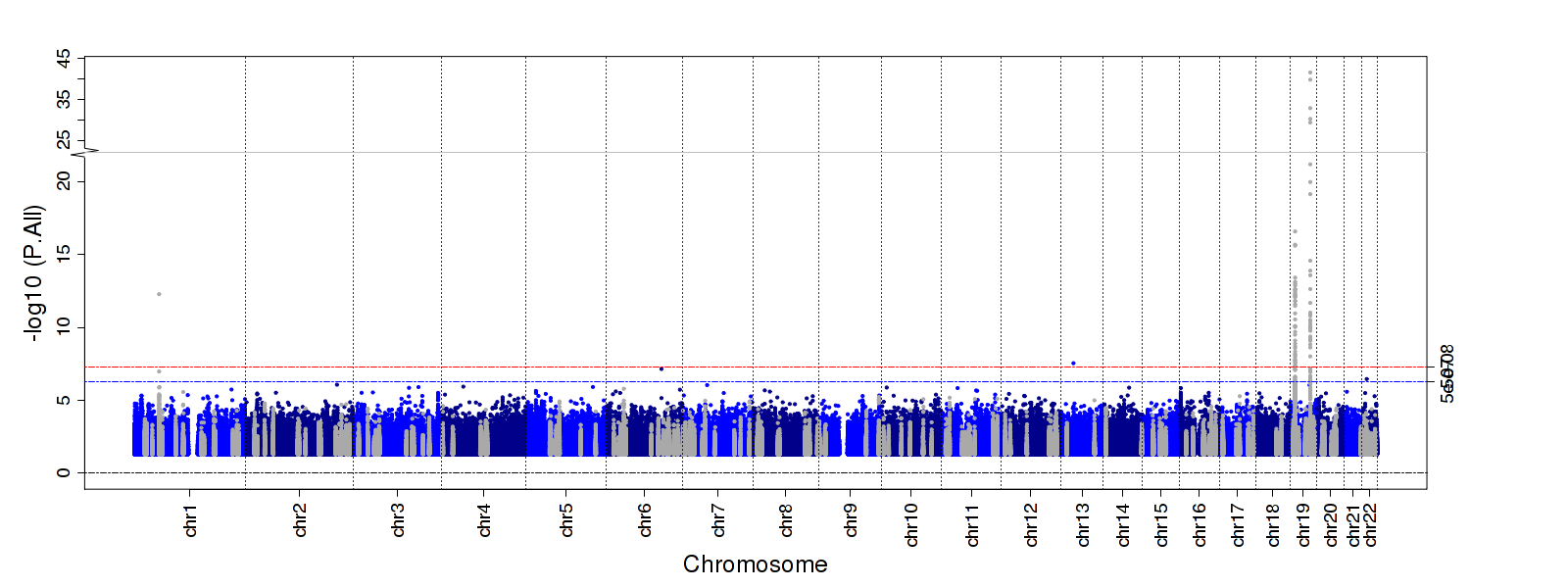

D. TG

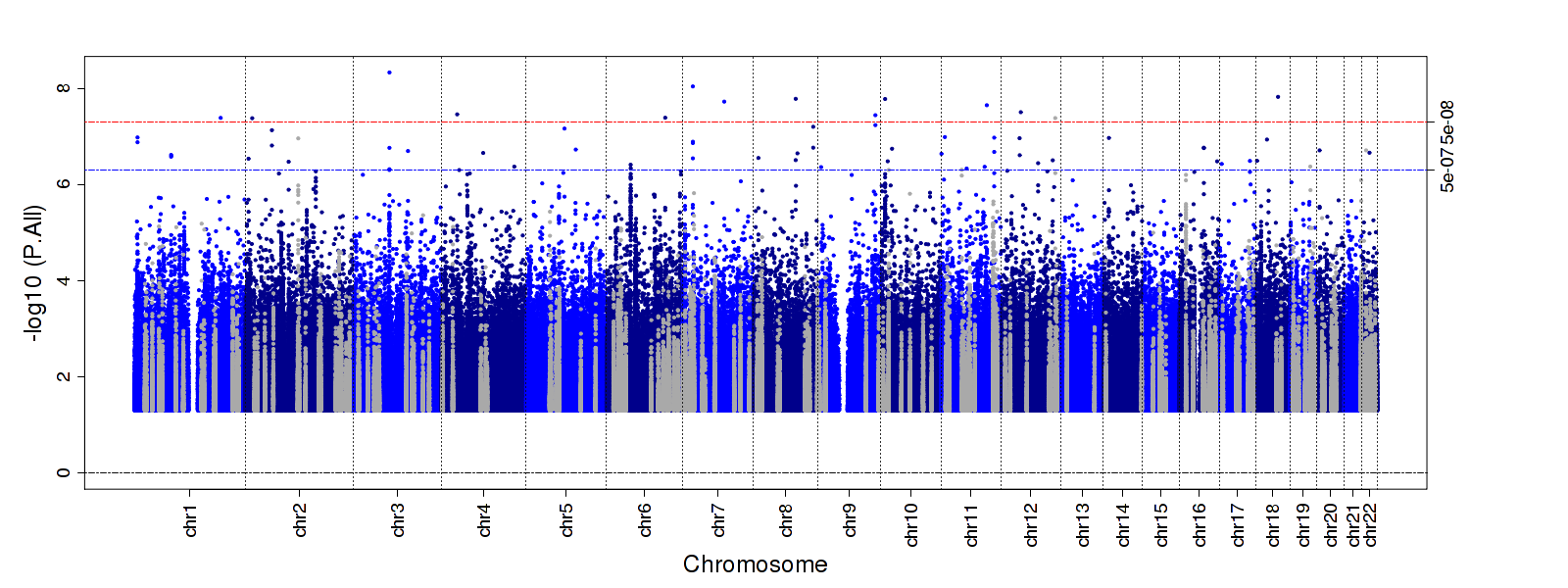

E. TG/HDL

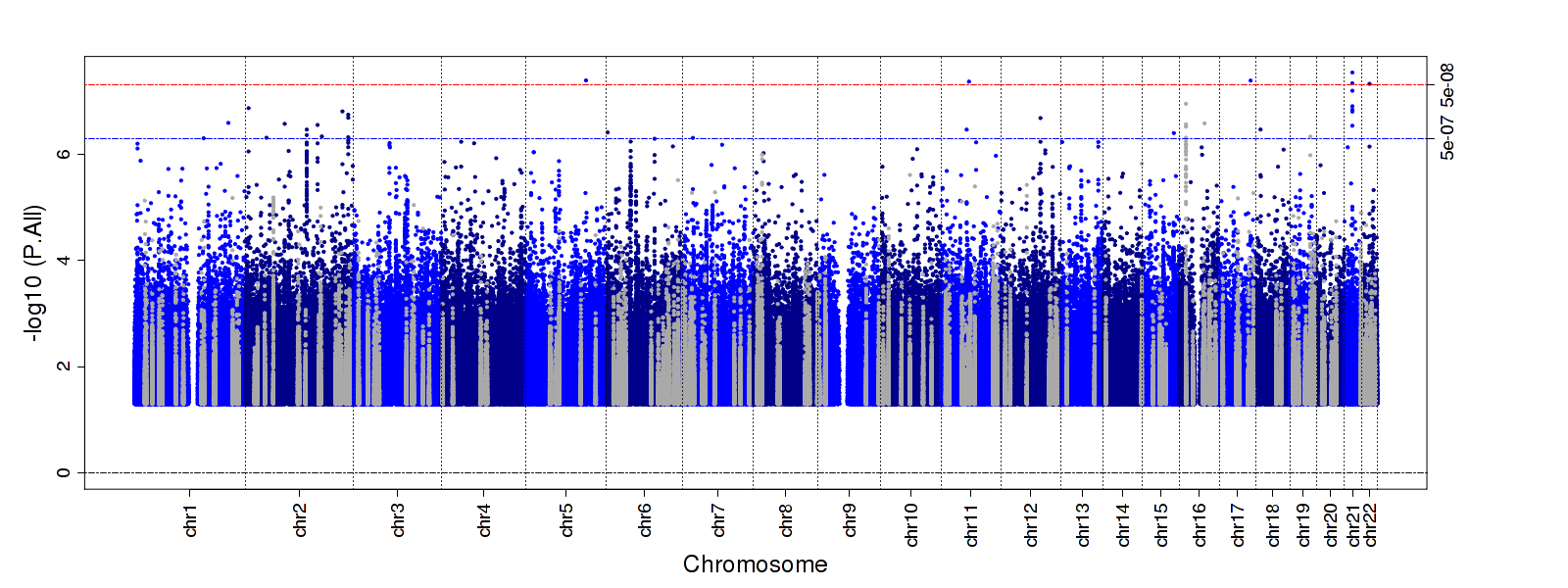

Supplementary Figure 2: QQ Plots for Serum Lipids in AADM in a model adjusted for age, age^2^, sex, and 3 PCs.

A. CHOL	B. HDL
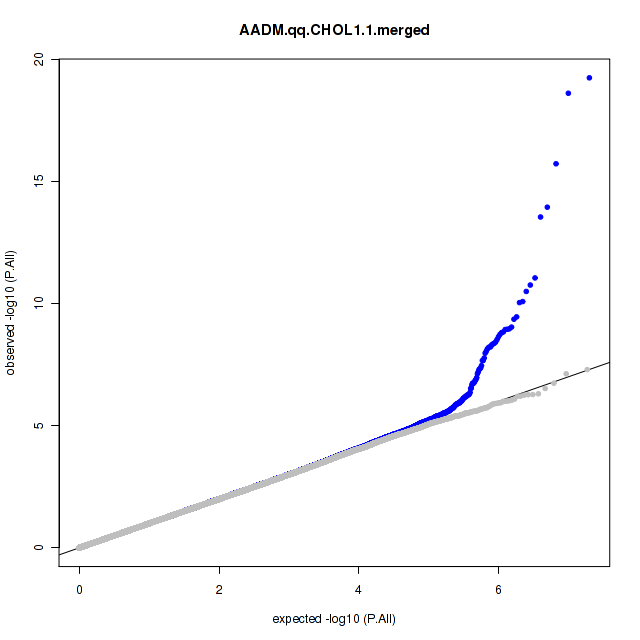

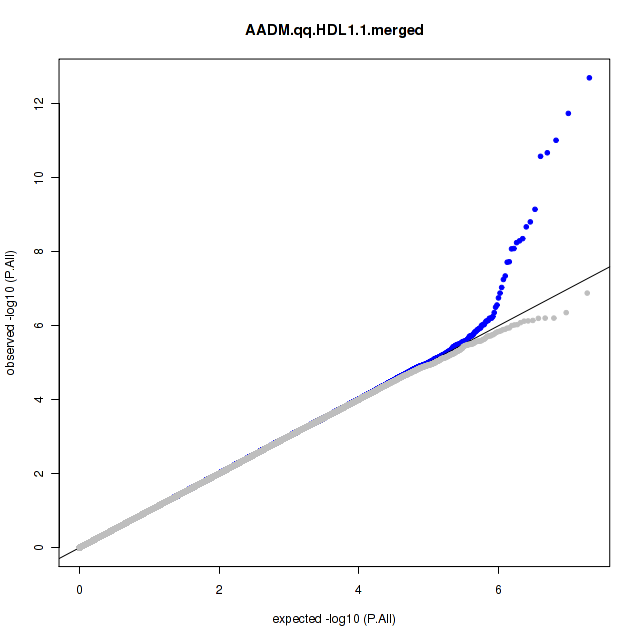

C.
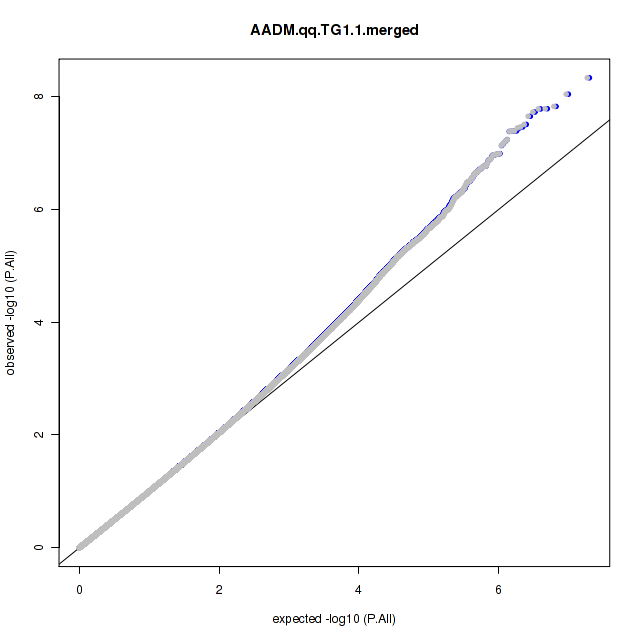

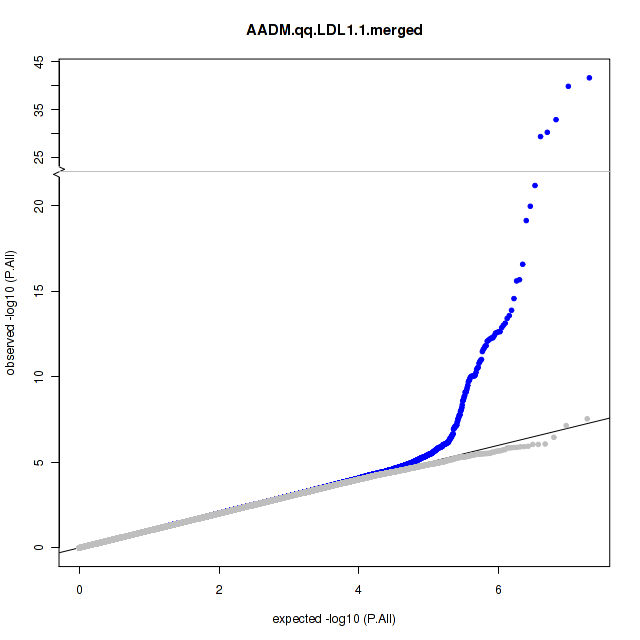
LDL	D. TG

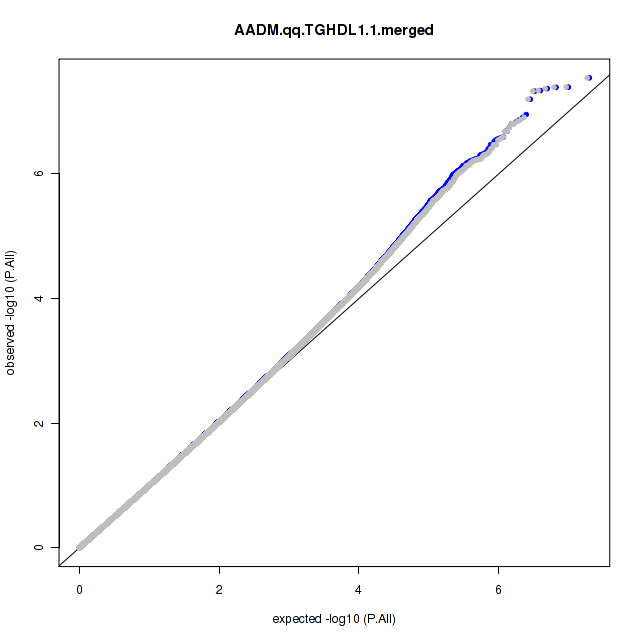
 E. TG/HDL
